# The swimming plus-maze test: a novel high-throughput model for assessment of anxiety-related behaviour in larval zebrafish. (*Danio rerio*)

**DOI:** 10.1101/342402

**Authors:** Zoltán K Varga, Áron Zsigmond, Diána Pejtsik, Máté Varga, Kornél Demeter, Éva Mikics, József Haller, Manó Aliczki

## Abstract

Larval zebrafish (*Danio rerio*) has the potential to supplement rodent models due to the availability of resource efficient methods implying high-throughput screening and high-resolution imaging techniques. Although behavioural models are available in larvae, only a few, insensitive approaches can be employed to assess anxiety. Here we present the swimming plus-maze (SPM) test paradigm to assess anxiety-related states in young zebrafish. The “+” shaped apparatus consists of arms of different depth representing differentially aversive context. The paradigm was validated i.) in larval and juvenile zebrafish, ii.) after administration of compounds affecting human anxiety and iii.) in differentially aversive experimental conditions. Furthermore, we compared the SPM with conventional “anxiety tests” of larvae such as the open tank and light/dark tank tests to identify their shared characteristics. We clarified that the preference towards deeper water is conserved trough the ontogenesis and can be abolished by anxiolytic or enhanced by anxiogenic agents, respectively. The behavioural read-out is insensitive to the aversiveness of the platform and unrelated to behaviours assessed by conventional tests utilizing larval fish. Taken together, we developed a sensitive high-throughput test measuring anxiety-related responses of larval zebrafish, which likely reflect bottom-dwelling behaviour of adults, potentially supporting larva-based integrative approaches.

## Introduction

Mental disorders, particularly those associated with anxiety represent a serious burden on the individual and the society as well (Wittchen and Jacobi, 2005). In order to resolve therapy of such disorders clinical and preclinical research makes efforts to uncover the basis and develop novel interventions in anxiety-related psychopathologies. However, compounds tested on the preclinical level despite the high resource intensity of this phase show low success rate in placebo controlled clinical trials (Haller and Aliczki, 2012; Haller et al., 2013). Consequently, there is an emerging need for innovative and resource-efficient approaches supporting the development of new therapeutic approaches.

Zebrafish (*Danio rerio*) is a tropical fish species (Parichy, 2015) rapidly gaining attention in biomedical research. The validity of this model is supported by the fact that 84% of human disease-related genes have at least one zebrafish orthologue (Howe et al., 2013). In addition, the central nervous system of teleost fish shares major characteristics with that of higher vertebrates, but possesses less complexity, which makes it very plausible to model basic brain functions (Broglio et al., 2005; Maximino et al., 2013; Panula et al., 2010; Stewart and Kalueff, 2014). Besides these homologies, the rationale for disease modelling in zebrafish is also supported by its low maintenance cost and reduced ethical concerns about its use (Lieschke and Currie, 2007; MacRae and Peterson, 2015; Stewart and Kalueff, 2014).

A particular advantage of zebrafish models is the availability of numerous manipulations and monitoring techniques of physiological processes in the larval stage. Due to the optical clarity of fish of this age, optogenetic manipulations and imaging of central nervous system activity is possible at the same time with a temporal and spatial resolution surpassing the rest of the models by far. Furthermore, larval fish show a wide behavioural repertoire, as early as a few days after hatching, making it an ideal model to combine behavioural and physiological screening. Despite that, there are relatively few approaches utilizing anxiety-related responses of larvae, which makes conducting such integrative phenomenological analysis difficult.

Furthermore, available behaviour tests utilizing larvae are based on the natural aversion of exposed (Richendrfer, 2012; Schnörr, 2012) or poorly lit areas (Bai et al., 2016; Cheng et al., 2016), but not water surface, albeit it is the most frequently measured and reproducible defensive response to novelty in adult zebrafish (Bencan and Levin, 2009; Blaser, 2010; Blaser and Rosemberg, 2012; Egan and Kalueff, 2009; Kysil et al., 2017; Levin, 2007; Maximino, 2010; Maximino et al., 2012; Sackerman, 2010; Stewart and Kalueff, 2011; Walsh-Monteiro, 2016). This is possibly due to the difficulties with recording the swimming depth of fish in a high-throughput context. Designing a behavioural paradigm in larvae assessing such typical defense response, particularly in combination with the wide range of techniques for modifying and monitoring physiological function would allow to assess the basis of anxiety-like states in a high-throughput and highly detailed manner.

In the current study, we present the swimming plus-maze (SPM) test, a paradigm for high-throughput screening of anxiety-related behaviour in larval zebrafish. The SPM platform consist of differentially deep arms, offering a choice between differently aversive zones, making high-throughput analysis of surface avoidance behaviour possible. The test is highly analogous to the rodent elevated plus-maze paradigm (Pellow et al., 1985; Walf and Frye, 2007), both in respect of its concept, design and observed pattern of behavioural outcome. To validate our test we i.) analyzed the effects of anxiolytic and anxiogenic compounds on larvae and juveniles, ii.) investigated the effects of experimental conditions such as light intensity, test repetition, and the context of test batteries. Furthermore, iii.) we aimed to determine the shared characteristics of SPM with previous conventional tests employed in larval zebrafish.

## Results

### Exploration pattern and correlation of variables in the SPM

The exploration pattern and the association of measured variables were calculated from the data of vehicle treated animals. Zebrafish prefer the deep over the shallow arms (est_zone:shallow_=-22.18, SE=3.83, df=45, t=-5.79, p=6.42e-07) and the center zone (est_zone:center_=-20.96, SE=3.83, df=45, t=-5.47, p=1.89e-06). The detailed exploration analysis and the heatmaps revealed an emerging trend in this preference towards the outermost parts of the deep arms (mean slope=4.59, SE=0.65) (Figure 1b-d). According to the correlational analysis there is a strong positive association between choice index and deep/total arm entries (cor_choice index:deep/total arm entries_=0.65, p = 2.003e-08), while both of these variables negatively correlate with the relative enter frequencies to the shallow arms (cor_choice index:shallow/total arm entries_=-0.79, p = 2.476e-14; cor_deep/total arm entries:shallow/total arm entries_ = −0.83, p = 2.272e-16). Neither of the latter variables (cor_choice index: velocity_ = −0.08, p = 0.522; cor_deep/total arm entries:velocity_= −0.14, p = 0.267; cor_shallow/total arm entries:velocity_ = −0.003, p = 0.982), but total arm entries is associated with velocity (cor_total arm entries:velocity_= 0.58, p = 1.019e-06) (Figure 1e).

**Figure 1.**
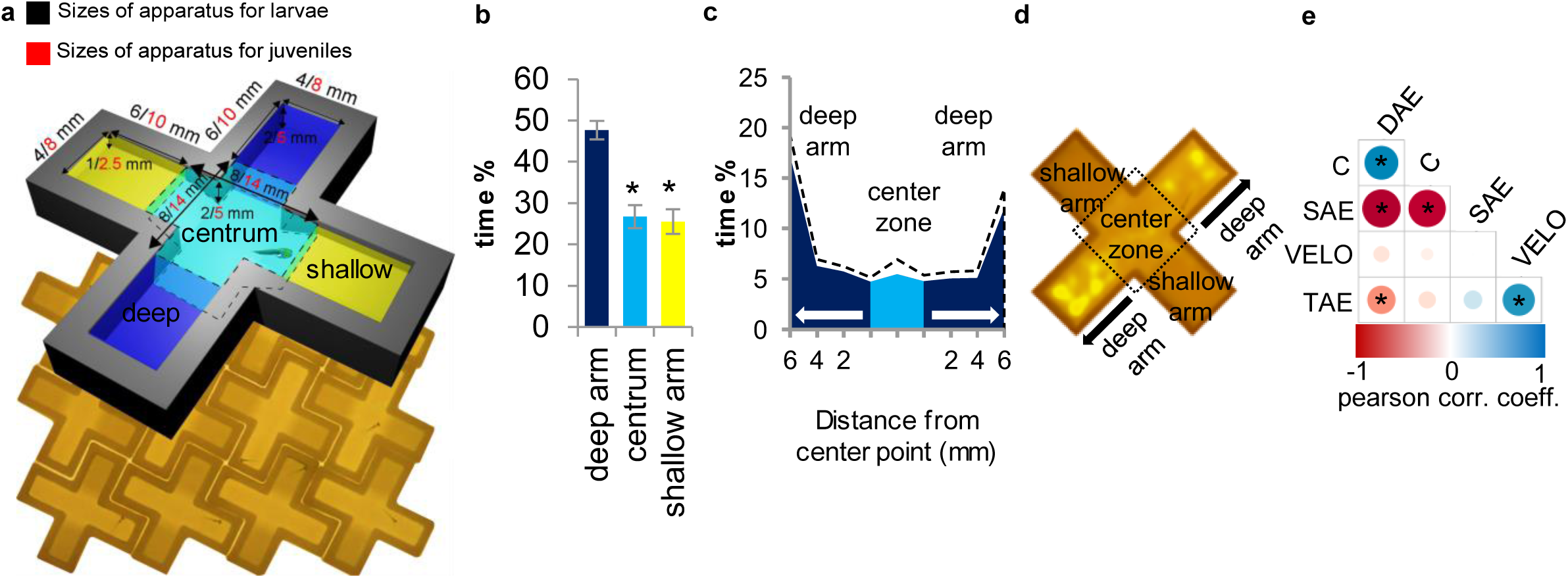
The swimming plus-maze test (SPM): apparatus, behaviour and selection of variables. Dimensions and spatial setting of the SPM platforms. Dashed lines indicate the boundary of areas (**a**). Spatio-temporal patterns of behaviour in the SPM: arm preference (**b**); higher resolution analysis (dashed lines indicate mean ± SEM) (**c**) and heatmap (**d**) of exploration pattern. *: significant differences from percentage of time spent in deep arms. Correlations between measured variables (**e**). DAE: deep/total arm entries, C: choice index, SAE: shallow/total arm entries, VELO: mean velocity, TAE: total arm entries. Size and color intensities of dots are in linear association with the correlation coefficients. *: significant correlation between two variables.

### Pharmacological validation of the SPM

In *Experiment 1a* we assessed the effects of the anxiolytic agent buspirone on larva behaviour. Vehicle treated fish has robust preference towards the deep arms over the shallow ones (est_zone:shallow_=-55.73, SE=19.77, df=30, t=-2.82, p=0.009) or the central zone (est_zone:center_=-66.63, SE=18.91, df= 26, t=-3.52, p=0.002). 50 mg/L buspirone decreased the time spent in deep arms (est_treatment:50mg/L_=-45.76, SE=15.66, df=67, t=-2.92, p=0.005) and, as the significant interaction indicates, anxiolytic treatment completely (est_treatment:50mg/L*zone:shallow_=58.72, SE=23.6, df=75, t=2.49, p=0.015; est_treatment:50mg/L*zone:center_=74.28, df=66, t=3.22, SE=23.07, p=0.002) or partially (est_treatment:100mg/L*zone:center_=50.22, SE=23.20, df=84, t=2.16, p=0.033) abolished the control ratio (Figure 2a). Buspirone did not affect choice index (est_treatment:25mg/L_=0.31, SE=0.36, df= 32, t= 0.88, p=0.358; est_treatment:50mg/L_=0.42, SE=0.37, df=32, t=1.12, p=0.182; est_treatment:100mg/L_=0.25, SE=0.38, df=31, t=0.68, p=0.5) (Figure 2b), however, significantly lowered deep/total arm entries (est_treatment:50mg/L_=-0.35, SE=0.15, df=31, t=-2.32, p=0.027) compared to control (Figure2c). None of the applied concentrations affected mean velocity (est_treatment:25mg/L_=0.69, SE=0.89, df=31, t=0.77, p=0.448; est_treatment:50mg/L_=-0.39, SE=0.90, df=29, t=-0.43, p=0.670; est_treatment:100mg/L_=-1.11, SE=0.92, df=30, t=-1.21, p=0.237) (Figure 2d).

**Figure 2.**
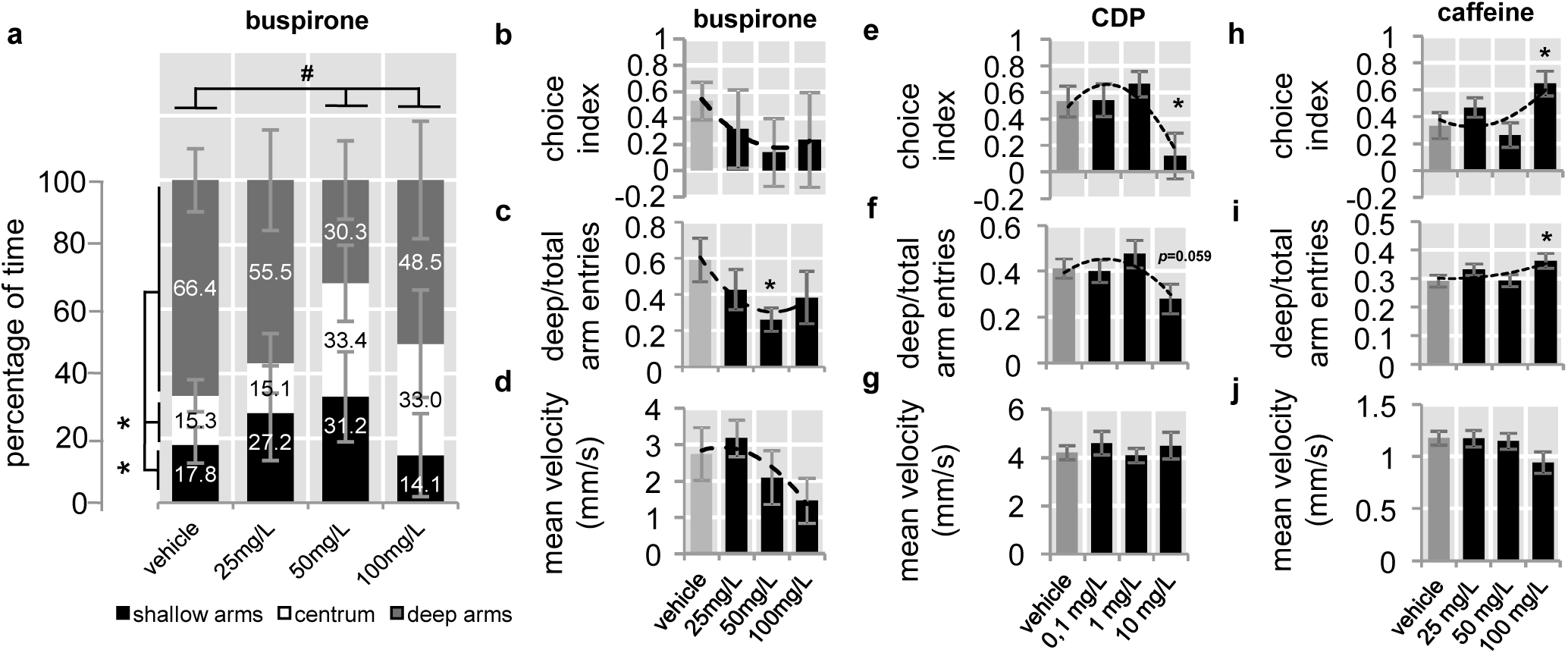
Pharmacological validation of the SPM. Changes in spatio-temporal pattern of behaviour in response to buspirone treatment (**a**). *: significant difference from time percentage spent in deep arms, #: significant interaction between treatment and area. Changes in arm preference associated variables in response to buspirone (**b-d**); chlordiazepoxide (CDP) (**e-g**); and caffeine (**h-j**). *: significant difference from vehicle treated group.

In *Experiment 1b* we investigated the effects of the anxiolytic agent CDP on larval behaviour. Vehicle treated fish showed highly positive choice index towards the deep arms, which was significantly lowered by 10 mg/L CDP (est_treatment:10mg/L_=-0.53, SE=0.18, df= 6.634e+01, t=3.38, p=0.024) (Figure 2e). CDP only marginally decreased deep/total arm entries (est_treatment:10mg/L_=-0.14, SE=0.07, df=64, t=-1.92, p=0.059) (Figure 2f) and did not affect mean velocity (est_treatment:10mg/L_=0.46, SE=0.52, df=63, 0.92, p=0.391) (Figure 2g).

In *Experiment 1c* we assessed the effects of the anxiogenic compound caffeine on larva behaviour. Control zebrafish showed strong deep arm activity, which was increased by the highest concentration of caffeine indicated by the enhanced choice index (est_treatment:100mg/L_=0.32, SE=0.13, df= 71.4, t=-2.40, p=0.019) (Figure 2h) and deep/total arm entries (est_treatment:100mg/L_= 0.07, SE=0.03, df= 70.1, t=-2.30, p=0.024) (Figure 2i). Also, caffeine had a marginal decreasing effect on velocity (est_treatment:100mg/L_=-0.22, SE=0.12, df= 74, t=-1.86, p=0.067) (Figure 2j).

### Validation of the SPM in juvenile zebrafish

In *Experiment 2*, we studied the validity of SPM in 30 dpf juvenile zebrafish by treatments with 50 mg/L buspirone, shown to be effective in the case of larvae, applying different incubation protocols (washout vs continuous treatment). Continuous exposure to buspirone significantly lowered positive choice index (est_treatment:continuous_=-0.74, SE=0.24, df=34, t=3.04, p=0.005) (Figure 3a) but not deep/total arm entries of fish (est_treatment:continuous_ =-0.04, SE=0.09, df=29, t=-0.47, p=0.646) (Figure 3b), however, both applied treatments decreased mean velocity (est_treatment:washout_=-2.42, SE=0.98, df=31, t=-2.46, p=0.020; est_treatment:continuous_= −33.92, SE=0.94, df=28, t=-4.15, p=0.0003) (Figure 3c).

**Figure 3.**
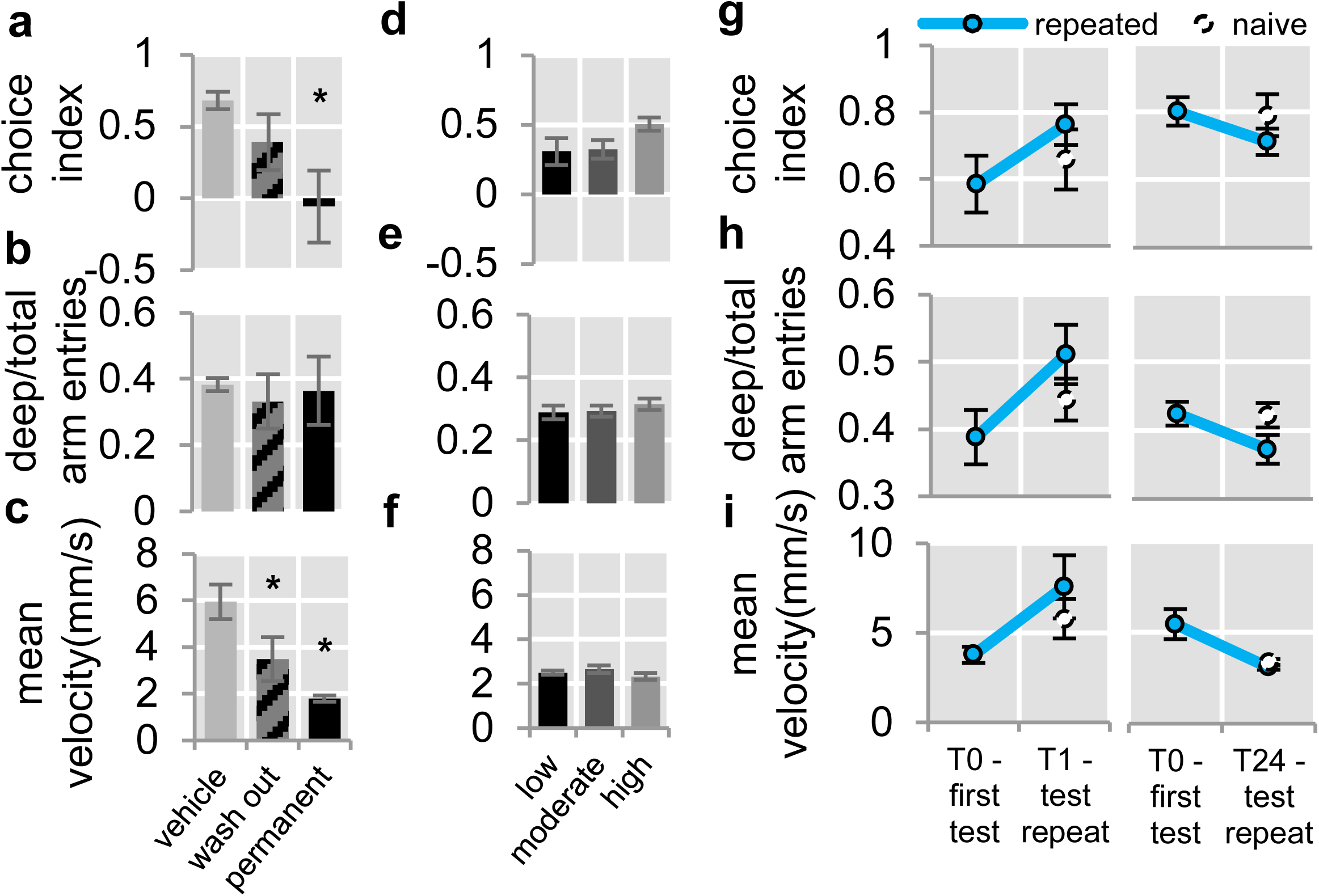
Validation in juveniles and the effect of environmental aversiveness and familiarity. Behaviour of juvenile zebrafish after buspirone treatment (**a-c**); Effects of different light intensities (**e-g**) and repetition of the test (**h-j**) on SPM behaviour. T0: results from the first test of repeated groups, T1 and T24: results from the tests of naive groups and the second tests of repeated groups after 1 and 24 hour intervals.

### Effects of environmental aversiveness on behaviour in the SPM

In *Experiment 3*, we analyzed the effects of environmental aversiveness on SPM behaviour by exposing juveniles to different light intensities. Results from the low light exposed group were set as the reference level of the model. Neither moderate nor intense illumination affected the measured behavioural variables compared to low light intensity group (for statistical data see Table S1) (Figure 3d-f).

### Effects of repeated testing in the SPM

In *Experiment 4a* and *4b* in order to assess the reproducibility of SPM, we tested juvenile zebrafish repeatedly in 1 and 24 hour intervals respectively, using additional naïve control animals in the second sampling. Results from the second test were set as the reference level of the model. Measured behaviour in the repeated test did not differ from their baseline values or from the results of naïve controls (Figure 3g-i) (for statistical data see Table S1).

### Shared characteristic of the SPM with conventional anxiety tests

In *Experiment 5* we compared behavioural data from SPM with conventional tests measuring anxiety-related responses of zebrafish. Juvenile zebrafish underwent the OT, SPM, and LDT tests in rapid succession. In order to assess whether behavioural outcomes of SPM are affected by prior OT testing, we introduced an additional parallel tested control group, naïve to any test exposure. Behaviour in the SPM was unaffected by previous OT exposure compared to test naïve controls (choice index: est_sampling:naïve=_ 0.12, SE=0.09, df=30, t=-1.31, p=0.202; deep/total arm entries: est_sampling:naïve_=-0.08, SE=0.07, df=30, t=-1.3, p=0.205) (Figure 4c-d). Surprisingly, mean velocity in the SPM and OT tests did not correlate significantly (r=0.32, p=0.241) (Figure 4b). Also, there were no significant correlation between choice indices of SPM and OT (r=0.07, p=0.794) or SPM and LD (r=0.47, p=0.076) (Figure 4e). However, according to the correlation coefficients between these variables, SPM and LD shared 22.9% of variance, in contrast to SPM-OT comparison in which this measure was only 0.49% (Figure 4f).

**Figure 4.**
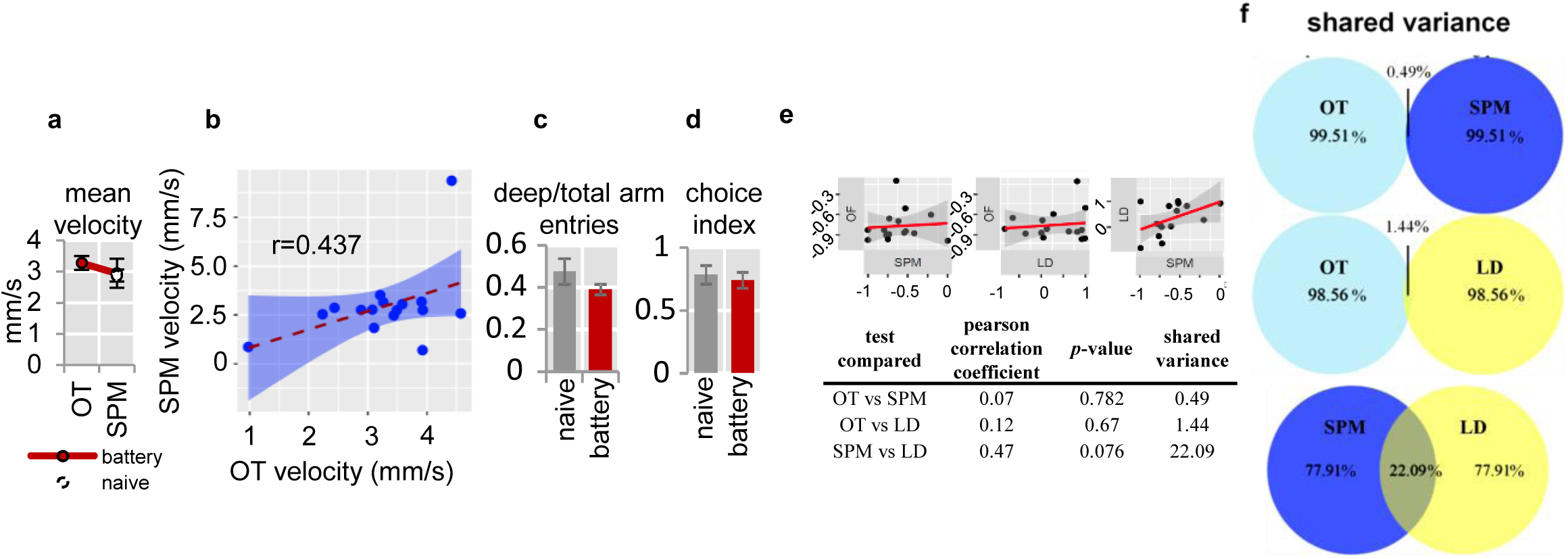
Shared characteristics of SPM and conventional “anxiety tests” of zebrafish. Values (**a**) and correlations (**b**) of mean velocity measured in the SPM and OT tests. Deep arm activity related variables measured in naïve and battery tested subjects **(c-d)**. Correlations between choice indices of “anxiety tests” of zebrafish (**e**). Shared variance between choice indices of tests shown by Venn diagrams (**f**).

## Discussion

Our results show that both larval and juvenile zebrafish prefer deep over shallow arms or the central zone. This preference was abolished by the clinical anxiolytics buspirone and CDP, while it was enhanced by the anxiogenic caffeine. SPM was applicable regardless of the developmental stage of the behavioural phenotype, the aversiveness or the familiarity of the testing environment, or prior OT exposure. Interestingly, deep arm activity in the SPM was unrelated to behavioural responses measured by conventional “anxiety tests” of larval zebrafish.

The primary behavioural responses in our study described arm preference and its background motivation. Choice index and deep/total arm entries are strongly correlated variables, however, neither of these are associated with the locomotion of zebrafish, suggesting that these variables are associated with closely related, if not the very same motivational states, hence potentially substitutable. This relationship indicates an important difference between SPM and EPM in which the relative entries to one of the closed arms, representing less aversive area, is rather a predictor of locomotion than anxiety (Walf and Frye, 2007). In the SPM the number of total arm entries correlates with velocity. However, we suggest to use the latter one due to its better resolution and the marked phenomenological differences between the movement of fish and rodents. In addition, we recommend to conduct such fine scale behavioural analysis in order to describe exploration patterns in detail.

Best to our knowledge, our group is the first that observed and utilized preference towards deeper water areas in larval zebrafish. Such marked preference of deeper zones possibly reflects the bottom-dwelling geotactic behaviour of zebrafish shown in response to novelty, described in adults by several groups using either the novel tank diving test or a plus maze with a ramp (Blaser and Rosemberg, 2012; Levin, 2007; Walsh-Monteiro, 2016). This parallelism is supported by the fact that in our experiments both larvae and post-metamorphic juveniles, showing completely developed behavioural phenotype (Dreosti et al., 2015; Lau and Guo, 2011), expressed similar behavioural pattern. There is a growing body of evidence in adult zebrafish that such phenomenon is rather driven by the aversion from water surface than preference for the bottom (Blaser and Goldsteinholm, 2012) and that direct exposure to such stimuli is highly stressful for fish (Fulcher et al., 2017; Piato et al., 2011). Furthermore, Ingebretson and Masino found that larval zebrafish show significantly less active swimming in shallower waterbodies, and also a higher ratio of zebrafish stayed completely immobile in such conditions (Ingebretson and Masino, 2013). This response possibly interpretable as an increase in anxiety-related freezing behaviour of larvae. The nature of this relationship between the relative depth of water and the level of aversiveness in larval stage is going to be the aim of future investigations. Interestingly, while the central zone of the SPM is equally deep as the deep arms, this area was less attractive to zebrafish in our study, suggesting that the intersection of the platform represents a different, possibly more aversive stimulus, e.g. higher environmental exposure. These results indicate that the exploration pattern in the SPM is driven by more than one stimuli and associated motivational states. Importantly, possible differences of light intensity in deep and shallow arms stemming from the different distance from the light source, are very unlikely to lead to deep arm preference, as exploration patterns were unaffected by different light conditions. This is also supported by the fact that larval and post-metamorphic fish showed very similar behaviour in the SPM, despite a switch occurring from light preferring to dark preferring phenotype between these developmental stages (Lau and Guo, 2011). Taken together, the observed behaviour is very likely the result of a trade-off between highly exposed aspects of the SPM, e.g. water surface or the intersection, and the innate urge of animals to explore the novel environment.

In our pharmacological validation experiments, deep arm activity was decreased by anxiolytics buspirone and CDP, while it was increased by the axiogenic caffeine, without either affecting locomotion. Since earlier reports on the effects of buspirone and benzodiazepines are rather inconsistent (Bohus et al., 1990; Collinson and Dawson, 1997; Gao and Cutler, 1993, 1993; Haller et al., 2004; Johnston and File, 1986), it is an important feature of the SPM that both agents, regardless of their target of action, affected behaviour in a similar manner. Furthermore, another advantage of the SPM is that, contrary to other tests (Schnörr, 2012), there was no measurable ceiling-effect potentially masking the anxiogenic properties of caffeine. Interestingly, the employed anxiolytics affected different, although strongly correlated, variables, pointing out the importance of detailed behavioural analysis. It is also important to note, that in contrast to earlier reports (Bencan et al., 2009), we did not detect the sedative effects of CDP, suggesting that the SPM potentially has lower sensitivity for measuring motor changes. Despite this small limitation, our findings suggest that the basis of behaviour shown in the SPM is anxiety-related.

Our experiment in juveniles revealed that the behavioural pattern of vehicle treated post-metamorphic zebrafish is very similar to that measured in larvae. The significance of applying such an experiment to 30 dpf fish is that most behavioural domains, e.g. social or light/dark avoidance behaviour, have already developed to mature form by this age (Dreosti et al., 2015; Lau and Guo, 2011). Based on our analysis, including fish from different developmental stages, and the investigation of adult behaviour by other labs (Blaser and Rosemberg, 2012; Levin, 2007; Walsh-Monteiro, 2016), it is very likely that surface avoidance behaviour remains conserved throughout the ontogenesis. Such phenomena enable the consistent use of SPM, regardless of developmental stage, over other current methods. However, 50 mg/L buspirone, previously shown to be effective in larvae, affected deep arm activity only in the case of the longer incubation protocol, but decreased locomotion in all treatment groups. This divergent behavioural pattern was probably rather due to the maturation of the CNS than differing ability to absorb the compound, since the treatment exerted sedative effect in every case. Revealing the basis of these surprising effects is going to be the aim of future investigations, however, our results suggest that SPM is applicable in different developmental stages, thus enabling it to be used for behavioural analysis during the early stages of ontogenesis.

In our experiments aiming to determine the effects of environmental conditions, neither different light intensities, nor repeated testing influenced behaviour. However, there is a non-significant trend in the test repetition experiment suggesting an enhanced excitation after the 1 hour and a mild habituation after the 24 hours intervals. These results suggest that these aspects of aversiveness or familiarity of the testing environment do not act as a strong biasing factors in the SPM, in contrast to its rodent analogue (File et al., 1993; Griebel et al., 1993; Pellow et al., 1985; Walf and Frye, 2007). Moreover, the use of naive controls revealed that there is no considerable fluctuation in the behavioural patterns in 1 or 24 h intervals. Our results suggest, that the SPM paradigm can be employed in a flexible manner even in a high-throughput context.

To determine whether deep arm activity in the SPM is related to other forms of avoidance behaviour measured by conventional “anxiety tests” of larval zebrafish, e.g. OT and LDT, we compared these. While the behavioural endpoints of different paradigms only slightly correlated, avoidance behaviour in the SPM and LD shared 22.9% variance, reflecting much more similarity between these zebrafish tests than between their rodent counterparts (Ramos et al., 2008). Despite this, the low correlation coefficients and the fluctuating basal locomotor activity between separate experiments highlight the shortcomings of using a single test. Since i.) OT exposure did not influence subsequent SPM behaviour, ii.) all three applied tests measure different phenomenological correlates of anxiety and iii.) are differentially sensitive to motor changes, we recommend the use of the applied test battery to analyze behavioural phenotypes in detail.

In summary, we determined that larval and juvenile zebrafish show a yet unobserved behavioural pattern, which likely reflects bottom-dwelling behaviour, exerted by complex stimuli of the SPM test. This pattern can be manipulated by pharmacological agents similarly as anxiety-based responses of higher vertebrates, including humans. With employment of the SPM paradigm we are able to screen genotypes, adverse experiences or any type of manipulations potentially affecting anxiety in a more detailed and highly efficient manner due to the utilization of larvae instead of adults. In addition, with the use of SPM, we are potentially able to combine state-of-the-art methods, based on the unique advantages of larvae, e.g. *in vivo* imaging techniques, with detailed behavioural analysis offered by the presented test battery.

## Materials and Methods

### Animals

Wild type (AB) fish lines were maintained in the animal facility of ELTE Eötvös Loránd University according to standard protocols (Westerfield, 2000). All protocols used in this study were approved by the Hungarian National Food Chain Safety Office (Permits Number: PEI/001/1458-10/2015 and PE/EA/2483-6/2016).Our subjects were unsexed animals aged 8 or 30 dpf (days post fertilization). Fish were group-housed and maintained in 13 h light/11 h dark cycle. Animals were terminated on ice immediately after each experiment.

### Drugs

All compounds were dissolved in E3 medium and administered as water bath followed by a brief washout (except when testing was conducted in treatment solution) in compartments of a 24-well plate. Each compartment (diameter=15.6 mm) contained 1.5 ml of treatment solution. We applied buspirone and caffeine at the concentrations of 0 (vehicle), 25, 50, and 100 mg/l, and chlordiazepoxide (CDP) at the concentrations of 0 (vehicle), 0.1, 1, and 10 mg/l. Concentrations were selected based on earlier studies (Bencan et al., 2009)(Egan et al., 2009). Compounds were obtained from Sigma-Aldrich.

### Behavioural test and analysis

#### The SPM test

The swimming plus-maze (SPM) apparatus is a “+” shaped platform consisted of 2 + 2 opposite arms, different in depth, connected by a center zone. The platforms were 3D printed by a Stratasys Objet30 printer using white opaque PolyJet resin. To test both larvae and juveniles we designed two types of apparatuses matching the sizes of the subjects (for specifications see Table 1 and Figure 1a). It is important to note that the center zone and deep arms had the same depth, however, exploration patterns (video S1) shown by the detailed exploration analysis (Figure 1c) and the heatmap of exploration (Figure 1d) suggest that individuals actively discriminate these areas. Shallow arms did not act as physical barriers of fish movement in either type of platforms (video S2).

**Table 1.**
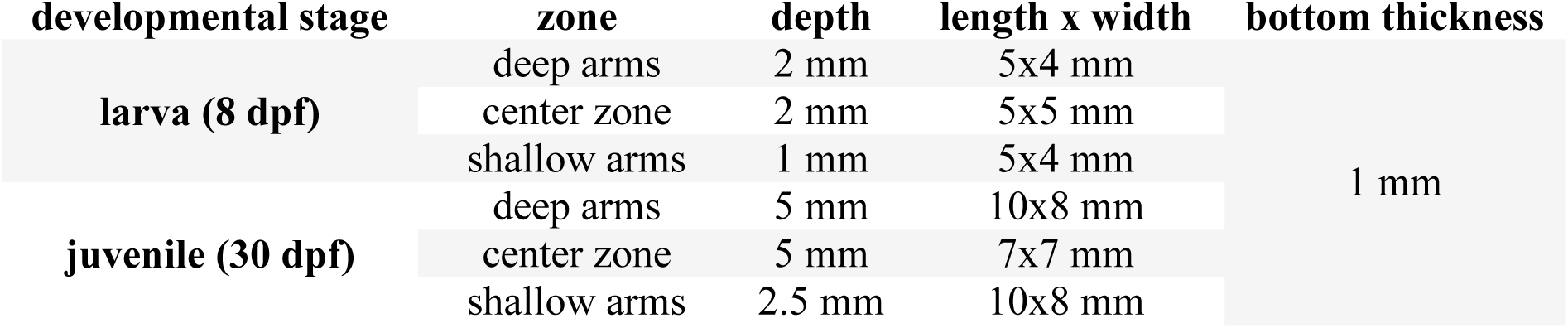

| developmental stage | zone | depth | length x width | bottom thickness |
| --- | --- | --- | --- | --- |
| larva (8 dpf) | deep arms | 2 mm | 5x4 mm | 1 mm |
|  | center zone | 2 mm | 5x5 mm |  |
|  | shallow arms | 1 mm | 5x4 mm |  |
| juvenile (30 dpf) | deep arms | 5 mm | 10x8 mm |  |
|  | center zone | 5 mm | 7x7 mm |  |
|  | shallow arms | 2.5 mm | 10x8 mm |  |

Experiments were conducted during the second part of the light phase, as zebrafish are reported to show continuous activity in this period (MacPhail, 2009). During a single trial, 8-16 subjects were tested simultaneously. The orientation of the platforms were randomized. Experiments in which pharmacological treatments were employed began with a 10 minute treatment bath in a 24-well plate compartment after which animals were gently pipetted to another compartment containing E3 medium for a brief washout (except when testing was conducted in treatment solution) then to the SPM for 5 or 10 minutes and recorded by a video camera. Experiments were carried out in an examination chamber and illuminated from beneath with white LED panels covered by matte Plexiglass. To reduce interfering stimuli from the environment the unit was covered with a black plastic box with a hole on top allowing the attachment of a video camera. Video recordings were analyzed with EthoVision XT 12 automated tracker software (NOLDUS L.P.J.J. et al., 2001).

Time spent in each zone and the mean of overall velocity was measured. To characterize arm preference, a choice index was defined as the relative time spent in the deep compared to in shallow arms. Consequently, choice index of 1 indicates 100% time in deep arms, whereas a choice index of −1 represents 100% time in the presumably more aversive shallow arms. Deep/total arm entries in SPM describes the relative frequency of the cases when an animal enters one of the deep arms. To conduct detailed exploration analysis of the deep arms and the center zone we measured the time spent in 3 equal complementary sections in each zones and calculated the average slope of these.

#### The OT test

For open tank (OT) testing, standard, completely transparent 24-well plates (d=15.6 mm) were used.

10 minutes long behavioural tests were conducted using the same protocol as in the case of SPM.

To describe thigmotaxis (edge preference) a choice index was defined as the relative time spent in the outer 20% of the compartment compared to time spent in the center zone. Mean of overall velocity was also measured.

#### The LDT test

For light/dark tank (LDT) testing, half of the compartments of a standard 24- well plate were masked with opaque matte black paint.

10 minutes long behavioural tests were conducted using the same protocol as in the case of OT and SPM.

To describe scotophobia (dark avoidance) a choice index was defined as the relative time spent in the dark of the compartment compared to time spent in the light zone.

### Experimental design

#### Pharmacological validation of the SPM

In *Experiment 1a*, *1b*, and *1c*, we validated the SPM paradigm by the administration of pharmacological agents shown to affect anxiety in preclinical rodent models and human clinical trials as well. Different sets of 8 dpf larvae were acutely treated with either the anxiolytics buspirone (0, 25, 50 and 100 mg/L) (*Experiment 1a*), CDP (0, 0.1, 1, and 10 mg/L) (*Experiment 1b*) or the anxiogenic caffeine (0, 25, 50 and 100 mg/L) (*Experiment 1c*) and their behaviour was recorded for 5 minutes. Sample sizes were 8-10 per group in *Experiment 1a*, 15-25 per group in *Experiment 1b*, and 15-26 per group in *Experiment 1c*.

#### Validation of the SPM in juvenile zebrafish

As juvenile, post-metamorphic fish avoid different stimuli than larvae, e.g. prefer dark over brightly lit environments (Lau and Guo, 2011), in *Experiment 2*, we determined the validity of the SPM paradigm for juvenile zebrafish. 30 dpf fish were acutely treated with 50 mg/L of buspirone the dose shown to be effective in larvae, and their behaviour was recorded for 5 minutes in the apparatus designed for juveniles. Since juvenile fish, in contrast to larvae are unable to absorb compounds through their whole body surface, we introduced an additional group which we placed into the testing apparatus in their treatment solution, providing longer incubation times (15 minutes). Sample sizes were 11-14 per group.

#### Effects of environmental aversiveness on behaviour in the SPM

In *Experiment 3* we assessed the effects of environmental aversiveness on behaviour in the SPM. 30 dpf juveniles were tested under different light intensities representing different levels of environmental aversiveness (Walf and Frye, 2007), and their behaviour was monitored for 10 minutes. As light conditions were set by the examination chamber lighting for all platforms in a particular trial, subjects from similar treatment groups were tested in one trial. The order of trials were randomized. Sample sizes were 16 animals in each group.

#### Effects of repeated testing in the SPM

In *Experiment 4*, we aimed to investigate the reproducibility of the SPM test. 30 dpf juvenile zebrafish were tested two times in 1 or 24 hour intervals and their behaviour was recorded for 10 minutes. In both cases, we tested a naïve control group as well in the second sampling period. Sample sizes were 10 in each group.

#### Shared characteristic of the SPM with conventional anxiety tests

In *Experiment 5*, we aimed to determine the shared characteristics of SPM with other tests measuring anxiety-like behaviour in larval zebrafish and its suitability for use in test batteries. 30 dpf juvenile zebrafish were tested for 10 minutes in the OT, SPM then the LDT test. In the case of SPM, we introduced an additional control group, without preliminary OT testing, to determine the effects of prior novelty exposure on SPM behaviour too.

### Data analysis

Data were represented as mean ± SEM. We performed statistical analysis using R Statistical Environment (R Core Team, 2017). To analyze the effects of pharmacological agents or experimental conditions on behavioural variables we used linear mixed models (Julian J. Faraway, 2016) from the *lme4* package (Douglas Bates, 2015). To separate variance stemming from time or sequence of experimental trials or location of test platforms, these factors were added as random effects to our models. To analyze within-group differences between percentage of time spent in each zone, we fitted linear mixed models with zone* treatment interaction as fixed and subject identifiers as random effects. We set treatment “vehicle” and zone “deep arms” as reference levels. We computed Pearson correlation coefficients (*r*) using *GGally* package (Barret Schloerke et al., 2017) and calculated percentage of shared variance applying the *r*^2^ × 100 formula. We calculated *p*-values from *t*-values of *lme4* using *lmerTest* package (Kuznetsova et al., 2016) and rejected H_0_ if *p*-values were lower than 0.05.

### Data availability statement

The datasets generated during the current study are available in the *figshare* repository (https://figshare.com/s/0ec47f2ffbe73f35500d).

## Compete and Interest

The authors declare no competing interest.

## Funding and Diclosure

This study was supported by the Hungarian Brain Research Program of the National Research, Development and Innovation Office (grant No 2017-1.2.1-NKP-2017-00002) and by the National Research, Development and Innovation Office grant No. 125390. to EM. MV is supported by the UNKP-17-4 New National Excellence Program of the Ministry of Human Capacities. MA is supported by the Janos Bolyai Research Scholarship of the Hungarian Academy of Sciences and by the National Talent Program Young Talents of the Nation Scholarship (NTP NTFO-17-C-0056). The authors declare no conflict of interest.

## Authors Contribution Statement

Conceptualization Z.K.V., M.A.; Methodology Z.K.V., M.A., A.Zs., M.V., K.D.; Formal Analysis Z.K.V.; Investigation Z.K.V., A.Zs., D.P.; Writing – Original Draft Z.K.V., M.A.; Writing – Review &amp M.A., E.M., M.V., A.Zs., K.D., D.P., J.H.; Editing Z.K.V., M.A.; Visualization Z.K.V.; Supervision M.A., E.M., M.V., J.H.

## Acknowledgement

We thank Gábor Wein (Lighting Application Specialist, Philips Lighting Hungary Kft., Budapest, Hungary) for the design and assembly of the applied examination chamber. We would also like to show our gratitude to Máté Tóth (Ph.D., IEM, HAS, Budapest, Hungary) for sharing comments that greatly improved the study.

## Supplementary material

**Table S1.**

| Experiment ID | compared groups | measure | estimate | SE | df | t-value | p-value |
| --- | --- | --- | --- | --- | --- | --- | --- |
| **3 (light intensities)** | low vs moderate light | choice index | 0.016 | 0.10 | 45 | -0.16 | 0.876 |
|  |  | deep/total arm entries | 0.004 | 0.03 | 45 | 0.15 | 0.879 |
|  |  | mean velocity | 0.16 | 0.19 | 38 | 0.84 | 0.408 |
|  | low vs intense light | choice index | 0.20 | 0.10 | 45 | -1.89 | 0.066 |
|  |  | deep/total arm entries | 0.03 | 0.03 | 45 | 0.94 | 0.355 |
|  |  | mean velocity | -0.18 | 0.19 | 38 | -0.95 | 0.351 |
| **4a (1 hour interval)** | baseline vs repeated | choice index | 0.19 | 0.14 | 6 | 1.31 | 0.239 |
|  |  | deep/total arm entries | 0.12 | 0.06 | 5 | -1.93 | 0.111 |
|  |  | mean velocity | 5.16 | 3.73 | 4 | -1.38 | 0.235 |
|  | repeated vs naïve | choice index | 0.15 | 0.11 | 26 | 1.36 | 0.184 |
|  |  | deep/total arm entries | 0.05 | 0.05 | 27 | -0.1 | 0.328 |
|  |  | mean velocity | 1.96 | 1.10 | 21 | -0.98 | 0.338 |
| **4b (24 hour interval)** | baseline vs repeated | choice index | 0.09 | 0.08 | 6 | -1.18 | 0.285 |
|  |  | deep/total arm entries | 0.05 | 0.03 | 8 | 1.64 | 0.141 |
|  |  | mean velocity | 2.46 | 1.62 | 6 | 1.52 | 0.179 |
|  | repeated vs naïve | choice index | 0.08 | 0.07 | 32 | -1.13 | 0.269 |
|  |  | deep/total arm entries | 0.05 | 0.03 | 32 | 1.95 | 0.060 |
|  |  | mean velocity | 0.34 | 0.28 | 33 | 1.19 | 0.241 |

## References

Bai, Y., Liu, H., Huang, B., Wagle, M., and Guo, S. (2016). Identification of environmental stressors and validation of light preference as a measure of anxiety in larval zebrafish. BMC Neurosci. 17.

Barret Schloerke, Jason Crowley, Di Cook, Francois Briatte, Moritz Marbach, Edwin Thoen, Amos Elberg, and Joseph Larmarange (2017). GGally: Extension to “ggplot2.”

Bencan, Z., and Levin, E.D. (2009). Buspirone, chlordiazepoxide and diazepam effects in a zebrafish model of anxiety. Pharmacol. Biochem. Behav. 75–80.

Bencan, Z., Sledge, D., and Levin, E.D. (2009). Buspirone, chlordiazepoxide and diazepam effects in a zebrafish model of anxiety. Pharmacol. Biochem. Behav. 94, 75–80.

Blaser, R.E. (2010). Behavioral measures of anxiety in zebrafish (Danio rerio). Behav. Brain Res. 56–62.

Blaser, R.E., and Goldsteinholm, K. (2012). Depth preference in zebrafish, Danio rerio: control by surface and substrate cues. Anim. Behav. 83, 953–959.

Blaser, R.E., and Rosemberg, D.B. (2012). Measures of Anxiety in Zebrafish (Danio rerio): Dissociation of Black/White Preference and Novel Tank Test. PLOS ONE 7, e36931.

Bohus, B., Koolhaas, J.M., Korte, S.M., Bouws, G.A., Eisenga, W., and Smit, J. (1990). Behavioural physiology of serotonergic and steroid-like anxiolytics as antistress drugs. Neurosci. Biobehav. Rev. 14, 529–534.

Broglio, C., Gómez, A., Durán, E., Ocaña, F.M., Jiménez-Moya, F., Rodríguez, F., and Salas, C. (2005). Hallmarks of a common forebrain vertebrate plan: specialized pallial areas for spatial, temporal and emotional memory in actinopterygian fish. Brain Res. Bull. 66, 277–281.

Cheng, R.-K., Krishnan, S., and Jesuthasan, S. (2016). Activation and inhibition of tph2 serotonergic neurons operate in tandem to influence larval zebrafish preference for light over darkness. Sci. Rep. 6.

Collinson, N., and Dawson, G.R. (1997). On the elevated plus-maze the anxiolytic-like effects of the 5-HT(1A) agonist, 8-OH-DPAT, but not the anxiogenic-like effects of the 5-HT(1A) partial agonist, buspirone, are blocked by the 5-HT1A antagonist, WAY 100635. Psychopharmacology (Berl.) 132, 35–43.

Douglas Bates (2015). Fitting Linear Mixed-Effects Models Using {lme4}. J. Stat. Softw. 67, 1–48.

Dreosti, E., Lopes, G., Kampff, A.R., and Wilson, S.W. (2015). Development of social behavior in young zebrafish. Front. Neural Circuits 9.

Egan, R.J., and Kalueff, A.V. (2009). Understanding behavioral and physiological phenotypes of stress and anxiety in zebrafish. Behav. Brain Res. 38–44.

Egan, R.J., Bergner, C.L., Hart, P.C., Cachat, J.M., Canavello, P.R., Elegante, M.F., Elkhayat, S.I., Bartels, B.K., Tien, A.K., Tien, D.H., et al. (2009). Understanding behavioral and physiological phenotypes of stress and anxiety in zebrafish. Behav. Brain Res. 205, 38–44.

File, S.E., Zangrossi, H., Viana, M., and Graeff, F.G. (1993). Trial 2 in the elevated plus-maze: a different form of fear? Psychopharmacology (Berl.) 111, 491–494.

Fulcher, N., Tran, S., Shams, S., Chatterjee, D., and Gerlai, R. (2017). Neurochemical and Behavioral Responses to Unpredictable Chronic Mild Stress Following Developmental Isolation: The Zebrafish as a Model for Major Depression. Zebrafish 14, 23–34.

Gao, B., and Cutler, M.G. (1993). Buspirone increases social investigation in pair-housed male mice; comparison with the effects of chlordiazepoxide. Neuropharmacology 32, 429–437.

Griebel, G., Moreau, J.-L., Jenck, F., Martin, J.R., and Misslin, R. (1993). Some critical determinants of the behaviour of rats in the elevated plus-maze. Behav. Processes 29, 37–47.

Haller, J., and Aliczki, M. (2012). Current animal models of anxiety, anxiety disorders, and anxiolytic drugs. Curr Opin Psychiatry.

Haller, J., Baranyi, J., Bakos, N., and Halász, J. (2004). Social instability in female rats: effects on anxiety and buspirone efficacy. Psychopharmacology (Berl.) 174, 197–202.

Haller, J., Aliczki, M., and Gyimesi Pelczer, K. (2013). Classical and novel approaches to the preclinical testing of anxiolytics: A critical evaluation.

Howe, K., Clark, M.D., Torroja, C.F., Torrance, J., Berthelot, C., Muffato, M., Collins, J.E., Humphray, S., McLaren, K., Matthews, L., et al. (2013). The zebrafish reference genome sequence and its relationship to the human genome. Nature 496, 498–503.

Ingebretson, J.J., and Masino, M.A. (2013). Quantification of locomotor activity in larval zebrafish: considerations for the design of high-throughput behavioral studies. Front. Neural Circuits 7.

Johnston, A.L., and File, S.E. (1986). 5-HT and anxiety: promises and pitfalls. Pharmacol. Biochem. Behav. 24, 1467–1470.

Julian J. Faraway (2016). Extending the Linear Model with R: Generalized Linear, Mixed Effects and Nonparametric Regression Models (CRC Press).

Kuznetsova, A., Per Bruun Brockhoff, and Rune Haubo Bojesen Christensen (2016). lmerTest: Tests in Linear Mixed Effects Models.

Kysil, E.V., Meshalkina, D.A., Frick, E.E., Echevarria, D.J., Rosemberg, D.B., Maximino, C., Lima, M.G., Abreu, M.S., Giacomini, A.C., Barcellos, L.J.G., et al. (2017). Comparative Analyses of Zebrafish Anxiety-Like Behavior Using Conflict-Based Novelty Tests. Zebrafish 14, 197–208.

Lau, B.Y.B., and Guo, S. (2011). Identification of a brain center whose activity discriminates a choice behavior in zebrafish. PNAS 108, 2581–2586.

Levin, E.D. (2007). Anxiolytic effects of nicotine in zebrafish. Physiol. Behav. 54–58.

Lieschke, P.D., and Currie, G.J. (2007). Animal models of human disease: zebrafish swim into view. Nat. Rev. Genet.

MacPhail, R.C. (2009). Locomotion in larval zebrafish: Influence of time of day, lighting and ethanol. Neurotoxicology 52–58.

MacRae, C.A., and Peterson, R.T. (2015). Zebrafish as tools for drug discovery. Nat. Rev. Drug Discov. 14, 721–731.

Maximino, C. (2010). Measuring anxiety in zebrafish: A critical review. Behav. Brain Res. 157–171.

Maximino, C., Benzecry, R., Oliveira, K.R.M., Batista, E. de J.O., Herculano, A.M., Rosemberg, D.B., Oliveira, D.L. de, and Blaser, R. (2012). A comparison of the light/dark and novel tank tests in zebrafish. Behaviour 149, 1099–1123.

Maximino, C., Lima, M.G., Oliveira, K.R.M., Batista, E. de J.O., and Herculano, A.M. (2013). “Limbic associative” and “autonomic” amygdala in teleosts: A review of the evidence. J. Chem. Neuroanat. 48–49, 1–13.

Noldus L.P.J.J., Spink A.J., and Tegelenbosch R.A.J. (2001). EthoVision: A versatile video tracking system for automation of behavioral experiments. Behav. Res. Methods Instrum. Comput. 3, 398–414.

Panula, P., Chen, Y.-C., Priyadarshini, M., Kudo, H., Semenova, S., Sundvik, M., and Sallinen, V. (2010). The comparative neuroanatomy and neurochemistry of zebrafish CNS systems of relevance to human neuropsychiatric diseases. Neurobiol. Dis. 40, 46–57.

Parichy, D.M. (2015). Advancing biology through a deeper understanding of zebrafish ecology and evolution. ELife 4.

Pellow, S., Chopin, P., File, S.E., and Briley, M. (1985). Validation of open:closed arm entries in an elevated plus-maze as a measure of anxiety in the rat. J. Neurosci. Methods 14, 149–167.

Piato, Â.L., Capiotti, K.M., Tamborski, A.R., Oses, J.P., Barcellos, L.J.G., Bogo, M.R., Lara, D.R., Vianna, M.R., and Bonan, C.D. (2011). Unpredictable chronic stress model in zebrafish (Danio rerio): Behavioral and physiological responses. Prog. Neuropsychopharmacol. Biol. Psychiatry 35, 561–567.

R Core Team (2017). R: A Language and Environment for Statistical Computing (Vienna, Austria: R Foundation for Statistical Computing).

Ramos, A., Pereira, E., Martins, G.C., Wehrmeister, T.D., and Izídio, G. S. (2008). Integrating the open field, elevated plus maze and light/dark box to assess different types of emotional behaviors in one single trial. Behav. Brain Res. 193, 277–288.

Richendrfer, H. (2012). On the edge: Pharmacological evidence for anxiety-related behavior in zebrafish larvae. Behav. Brain Res. 99–106.

Sackerman, J. (2010). Zebrafish Behavior in Novel Environments: Effects of Acute Exposure to Anxiolytic Compounds and Choice of Danio rerio Line. Int. J. Comp. Psychol. 23, 43–61.

Schnörr, S.J. (2012). Measuring thigmotaxis in larval zebrafish. Behav. Brain Res. 367–374.

Stewart, A., and Kalueff, A.V. (2011). Pharmacological modulation of anxiety-like phenotypes in adult zebrafish behavioral models. Prog. Neuropsychopharmacol. Biol. Psychiatry 1421–1431.

Stewart, A., and Kalueff, A.V. (2014). Zebrafish models for translational neuroscience research: from tank to bedside. Trends Neurosci. 37.

Walf, A.A., and Frye, C.A. (2007). The use of the elevated plus maze as an assay of anxiety-related behavior in rodents. Nat. Protoc. 2, 322–328.

Walsh-Monteiro, A. (2016). A new anxiety test for zebrafish: Plus maze with ramp. Psychol. Neurosci. 9., 457–464.

Wittchen, H.-U., and Jacobi, F. (2005). Size and burden of mental disorders in Europe—a critical review and appraisal of 27 studies. Eur. Neuropsychopharmacol. 15, 357–376.

